# Insight into the Interaction between Neuronal Calcium Sensors and Insulin

**DOI:** 10.1101/2024.07.26.605271

**Authors:** Uday Kiran, Venu Sankeshi, Yogendra Sharma

## Abstract

Calcium is crucial in insulin biology and Ca^2+^ sensor proteins enforced in insulin release and signalling. The neuronal calcium sensor proteins (NCS) such as NCS-1 and VILIP are shown to be involved in insulin secretion from β-pancreatic cells. However, the expression of different NCS proteins in the pancreas and their functional significance and role in pathologies remained unexplored. The present work, through different biophysical methods, presented that NCS proteins interact with insulin. NCS-1, the founder member of NCS family proteins interacts with insulin in a Ca^2+^ independent manner and Ca^2+^ enhances the affinity of the interaction. The evolutionarily conserved cryptic EF-hand in NCS proteins was found to be an essential commodity for binding insulin. The presence of Ca^2+^ binding first EF-hand abolishes the interaction with insulin and suggests the significance of non-functional EF-hand. The fluorescence and circular dichroism (CD) spectroscopy show that insulin interaction induces structural changes in NCS-1, which is demonstrated by size exclusion chromatography and analytical ultra-centrifugation. The autism mutant NCS-1-R102Q relatively retained insulin binding properties but with a significant difference in binding thermodynamics. Considering substantial sequence similarity among different NCS proteins and localisation in the pancreas, we examined the insulin interaction with the neurocalcin delta (NCALD). The NCALD shows metal ion-independent insulin binding and contrary to NCS-1, the Ca^2+^ abolishes the insulin binding. This highlights the differential regulation of Ca^2+^ towards insulin interaction in NCS protein. Conclusively the present work highlight that NCS proteins interact with insulin and further investigation would aid to understand the significance of NCS proteins in insulin physiology/pathophysiology and possible new molecular targets in diabetes.

## Introduction

Irrespective of cell type, **t**he Ca^2+^ plays a versatile and vital role in different cellular functions via Ca^2+^ binding/sensor proteins. The neuronal calcium sensor proteins (NCS) form the major class of Ca^2+^ sensors, involved in multiple cellular functions. NCS-1 is a founding member of the NCS family of proteins first identified in drosophila and highly conserved from yeast to humans [1]. The neuronal tissues show a high expression of NCS-1, however endocrine, heart, and gastrointestinal tract tissues also display NCS-1 [2, 3]. The significance of NCS-1 in the neuronal framework has been broadly studied and shown to be involved in neuronal plasticity, growth, regeneration, memory, exocytosis, and neurotransmission [1, 4]. The expressional deviance, mutations, and misfolding of NCS-1 are associated with bipolar disorder & schizophrenia, depression, autistic spectrum disorder (ASD), Parkinson’s disease, and neurodegeneration [4]. Along with NCS-1, other NCS family proteins such as hippocalcin, NCALD, VILIP, GCAPs, and KChIPs expressed in multiple tissues and genetic alterations demonstrated different pathological outcomes by deregulation of Ca^2+^ signalling [1]. In line with this, the non-neuronal pathological implications of NCS proteins and Ca^2+^ sensors gained attention since the first report on the significance of Ca^2+^ in insulin secretion [5]. The Ca^2+^ role in insulin biology has been explored extensively [6]. Deviation in mitochondrial Ca^2+^ homeostasis negatively affects insulin release, where the knock-out of Ca^2+^ sensor protein mitochondrial Ca^2+^ uptake 1 (MICU1) results in disruption of mitochondrial Ca^2+^ homeostasis [7]. Wolfram syndrome results in diabetes and optic atrophy caused by defective communication between the endoplasmic reticulum (ER) and mitochondria due to loss of function of ER-protein WFS1. NCS-1 interacts with WFS1 and assists its normal function, however, in wolfram syndrome NCS-1 abundance is low and retention of NCS-1 in WFS1defective cells re-established the ER and mitochondrial communication [8].

In the pancreatic context, NCS-1 and VILIP are shown to express and co-localise with insulin and potentiate insulin secretion via different pathways [9, 10]. NCS-1 increases exocytosis by priming secretory granules along with increasing the granules in the readily releasable pool through the interaction with Phosphatidylinositol 4-Kinaseβ (PIKβ) [9]. Over-expression of VILIP-1 enhanced insulin secretion in a cAMP-dependent manner and down-regulation was accompanied by decreased cAMP but increased insulin gene transcription [10]. NCS-1 knockout (KO) mice show anxiety and depressive behavior, however, recent study highlights that NCS-1 deficient mice develop obesity and diabetes by adulthood [11, 12]. Under high-fat diet (HFD), the NCS-1-KO mice are hyperglycemic and hyperinsulinemic suggests insulin resistance. Interestingly NCS-1 is associated with insulin receptor (IR) regulation and translocation in adipocytes and NCS-1 co-immunoprecipitated with IR complex [12]. The insulin-stimulated translocation of GLUT-4 to the plasma membrane in adipocytes, is negatively regulated by NCS-1 via PI 4-kinase interaction [13]. The insulin-stimulated translocation of GLUT-4 is associated with glucose clearance from plasma and defects are associated with insulin resistance and diabetes [14, 15]. Along with NCS-1, other NCS proteins are also implicated in insulin regulation and diabetes [16, 17]. Downstream regulatory element (DRE) antagonist modulator (DREAM) binds to DRE of insulin gene transcription promotor as dimmer under low Ca^2+^ levels and released at high Ca^2+^ levels inside the nucleus [16]. Polymorphism in 3’UTR in the NCALD gene affects the mRNA stability and results in susceptibility to diabetic nephropathy [17]. NCS proteins are drug target for insulinotropic (which potentiates the insulin release) agent repaglinide, which binds to NCS proteins and modulates their activity [18]. Considering the significant role of NCS proteins in diabetes, in this work, we attempted to find the direct interaction of insulin with NCS proteins. Here we present that NCS proteins bind to insulin and Ca^2+^ play role in the interaction. Further, we show the functional significance of cryptic EF-hand in NCS proteins and the consequence of pathogenic NCS-1 mutant on insulin binding.

## Materials and Methods

### Protein Overexpression and Purification

Different NCS genes from rat cDNA were cloned into pET21a vector. The recombinant proteins were over-expressed in *E*. coli BL-21 (DE3) cells with 0.5 mM IPTG induction. All the proteins were expressed in the soluble fraction. The NCS-1 and mutant overexpression, myristoylation, and purification were done as previously described [19]. In the case of NCALD and VILIP, the induced cells were incubated at 18 ºC and 37 ºC for 8 and 4 hours respectively. Both the proteins were purified using a Q-sepharose column (pH-7 for NCALD and pH-8 for VILIP). The cells were sonicated in lysis buffer (50 mM Tris, 1 mM EDTA, deoxycholic acid, Dnase, and PMSF) and the centrifuged supernatant was loaded on to column followed by a wash with 10 column volumes of wash buffer (50 mM Tris and 1 mM EDTA). The elution was achieved by gradient increment of NaCl concentration up to 500 mM. The eluted samples were concentrated and buffer exchanged with 50 mM Tris, pH-7.5, and KCl 100 mM buffer, followed by size exclusion chromatography using Sephadex G-75 (GE Healthcare) column. The pure fractions were incubated with a 10x molar concentration of EDTA for 1 hour and followed by thorough buffer exchange with Chelex-treated buffer (50 mM Tris, pH-7.5, and KCl 100 mM) to prepare decalcified protein fractions. The insulin (human) stocks were prepared by dissolving the powder (PAN biotech, Cell biology grade) in a buffer with 50 mM glycine, 100 mM KCl, and pH 2.5. The experimental concentrations were achieved by diluting the stock in 50 mM Tris, pH-7.5, and KCl 100 mM buffer.

### Site-Directed Mutagenesis

The NCS-1 R102Q mutant was generated by using Q5^®^ Site-Directed Mutagenesis Kit. The WT-NCS-1 plasmid was subjected to amplification with mutagenic oligonucleotides by following the manufacturer’s protocol. The amplified PCR product was subjected to DpnI digestion followed by transformation. The mutation was confirmed by an in-house sequencing facility.

### Fluorescence Spectroscopy

Hitachi F-7000 spectrofluorometer (Hitachi Inc., Japan) was used to record the intrinsic and extrinsic fluorescence. For Trp fluorescence experiments, 5 µM respective protein was taken in Chelex-purified 50 mM Tris, pH 7.5, and 100 mM KCl buffer, and insulin titrations were done. The quartz cuvette of the path length of 4 mm was used to record fluorescence spectra from 310–450 nm, at an excitation wavelength of 295 nm, a scan speed of 240 nm/min, and slits of 5 nm open on both sides. In the case of ANS fluorescence experiments, 5 μM protein was incubated with 100 μM of ANS (prepared 10 mM stock solution in dimethyl sulfoxide/water mixture), and emission spectra were recorded between 400-600 nm at an excitation wavelength of 370 nm. All the spectra were corrected for respective insulin blank and ANS-buffer blank.

### Circular Dichroism (CD) Spectroscopy

The Jasco-810 spectropolarimeter was used to record near- and far-UV CD spectra. A protein solution of 30 µM in 50 mM Tris, pH 7.5, and 100 mM KCl, was used for near-UV CD spectra record between 350-250 nm with a quartz cuvette of 1 cm path length and averaged with 4-6 accumulations. Similarly, using a quartz cuvette of 0.1 cm path length and with ∼ 5 µM protein solution, all far-UV CD spectra were recorded between 250-200 nm with 4 accumulations. All the titrations were corrected with respective buffer blanks in the Chelex-treated buffer.

### Analytical gel filtration

The NCS-1, insulin, and their complexes (∼ 90 µM) in apo and holo conditions were resolved on Superdex 75 (10/300) column (Wipro GE Healthcare) and absorption was recorded at 280 nm. The column was pre-equilibrated with buffer (50 mM Tris, pH 7.5, 100 mM KCl) consisting of either 0.1 mM EDTA or 2 mM Ca^2+^.

### Analytical ultracentrifugation (AUC)

AUC was carried out on a Beckman Optima XL-I Analytical Ultracentrifuge (Beckman Coulter, Fullerton, CA, USA) operating in sedimentation velocity mode with an aluminum centerpiece. Protein samples (∼ 90 µM) were prepared in 50 mM Tris, pH 7.5, 100 mM KCl buffer and centrifuged at a speed of 42,000 rpm at 25 °C in a Ti60 rotor, and data was collected using 280 nm absorbance optics for a total of 60 scans with the rate of one scan per every 5 minutes. The data were analysed using the c(s) method in SedFit software.

### Isothermal Titration Calorimetry

The metal ion binding and insulin binding for all proteins and mutants were performed on Microcal, VP-ITC. Decalcified protein solution of 30 μM concentration was prepared in Chelex-purified 50 mM Tris, pH 7.5, and 100 mM KCl buffer for metal ion binding studies. For insulin binding studies protein concentration of 300 μM was taken in a syringe and a 30 μM insulin solution was taken in the cell. The titrations were performed with a 4 μl injection volume with a spacing of 240 seconds until saturation was achieved. All titrations were corrected with respective buffer blanks.

### Insulin amyloidogenesis

The insulin fibrillation experiments were performed by taking 60 µM insulin alone or insulin and NCS-1 mixture in 50 mM glycine, pH 2.5, 100 mM KCl, with 10 µM ThT. The samples were incubated at 55 ºC (in the absence of Ca^2+^) or 60 ºC (in the presence of Ca^2+^) on a temperature-controlled orbital shaker at 500 rpm [20]. The samples were checked for a change in ThT fluorescence with an interval of 30 minutes using a 96-well plate on a Perkin-Elmer multimode plate reader. Readings were taken in triplicates for each sample and standard error was calculated for plotting.

## Results and Discussion

### Insulin induces changes in the Trp environment of NCS-1

Based on co-localisation with insulin and being a protein of wide interactome and a recent study on secretagogin a Ca^2+^ sensor binds insulin, we conceived the idea that NCS-1 could interact with insulin. By taking advantage of insulin being a tryptophan-less peptide hormone, we examined the insulin-induced change in tryptophan fluorescence of NCS-1. The tryptophan (Trp) fluorescence is extremely sensitive to the polarity of the environment and the change in Trp fluorescence reveals the local structural changes of protein [21]. NCS-1 contains two tryptophans, one in the N-terminal with one Ca^2+^ binding EF-hand and another in the Ca^2+-^ specific EF-hand containing the C-terminal domain [22]. We performed titration of insulin with NCS-1 in the presence (holo) and absence (apo) of Ca^2+^. In apo condition, the NCS-1 fluorescence shows λ max of ∼343 nm, and insulin titration resulted in a slight increase in fluorescence intensity (Fig. 1A). The addition of Ca^2+^ resulted in a significant increase in fluorescence of NCS-1 and a ∼2 nm blue shift in λ max suggests the change in the Trp environment (Fig. 1B). Titration with insulin in holo condition resulted in the movement of Trps into a more non-polar environment indicated by a blue shift of ∼2 nm in λ max (Fig 1B). The fluorescence results indicate the change in the Trps environment in NCS-1 protein due to insulin interaction.

**Fig. 1.**
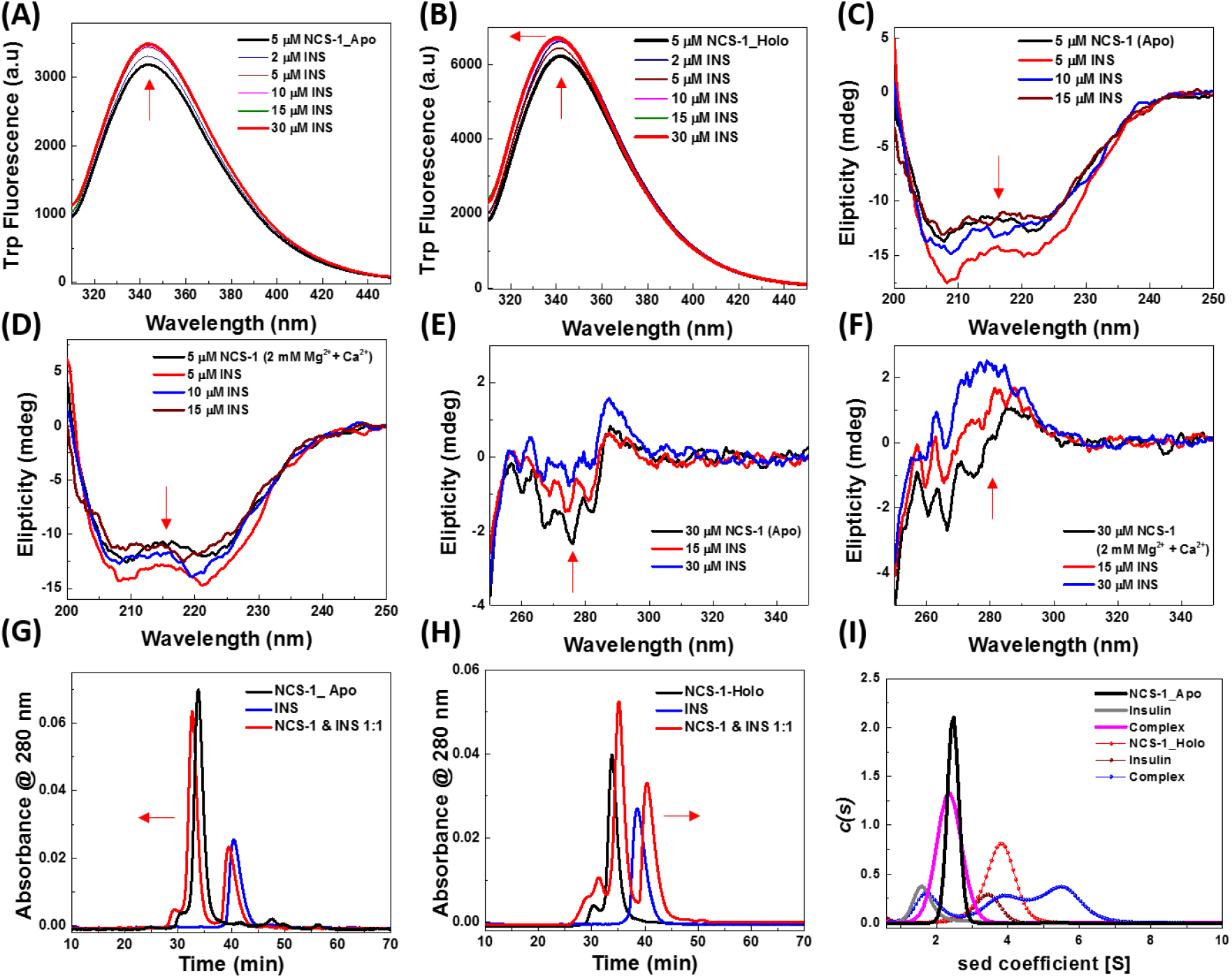
Insulin interaction induces structural changes in NCS-1 and forms a complex. (A) Apo (B) 2 mM Mg^2+^ & Ca^2+^ (holo) change in NCS-1 Trp fluorescence by insulin titration, NCS-1 (Black), and insulin saturated spectra (Red). (C) Apo (D) Holo, far-UV CD shows a change in secondary structural features of NCS-1 by insulin titration, NCS-1 (Black), 5 (Red), 10 (Blue), 15 (Wine) µM insulin. (E) Apo (F) Holo, near-UV CD spectra of NCS-1 by insulin titration, NCS-1 (Black), 15 (Red), 30 (Blue) µM insulin. (G) Apo (H) Holo Differential mobility of NCS-1, inulin, and complex checked by analytical gel filtration, NCS-1 (Black), insulin (Blue), and complex (Red). (I) The sedimentation coefficient of NCS-1, insulin, and complex was observed by analytical ultra-centrifugation. Apo, NCS-1 (Black), insulin (Gray) complex (Magenta). Holo, NCS-1 (red), insulin (Wine), and complex (Blue).

### Insulin induces structural changes in NCS-1

The secondary and tertiary structural elements of protein can be studied by using circular dichroism. Further, the change in spectral properties also helps to understand the protein-protein interactions [23]. The far-UV CD spectra indicate typical helical secondary structural features of NCS-1. The insulin titration resulted in a gain in negative ellipticity and a change in minima near 210 nm (Fig. 1C). Similar observations were made in the presence of 2 mM Ca^2+^ and maximum gain in ellipticity was prominent in the 1:1 ratio of NCS-1 and INS (Fig.1D). The results indicate the alteration in secondary structural elements of NCS-1 due to insulin.

The near UV CD spectra display changes in the Trp environment (Fig 1E). The spectral changes were further prominent in the presence of 2 mM Ca^2+^ (Fig. 1F). The change in ellipticity (towards the positive side) was observed near 270-280 nm wavelength in both conditions. From this, it is clear that insulin interaction with NCS-1 leads to substantial structural changes.

### Solution properties of NCS-1 and insulin complex

To understand the solution properties of NCS-1 and insulin complex, analytical gel filtration (AGF) and analytical ultra-centrifugation (AUC) were employed. The NCS-1 (black line) and insulin (blue line) were resolved on AGF individually and as a 1:1 molar ratio complex (red line) in apo (100 µM EDTA) and holo (2 mM Ca^2+^) conditions. In apo condition, the complex was eluted earlier compared to NCS-1 and insulin alone, suggesting the presence of higher molecular weight species (Fig. 1G). Contrary to apo, in the presence of 2 mM Ca^2+^, the complex eluted later compared to individual proteins (Fig. 1H). The results suggest the differential complex conformations of NCS-and insulin in apo and holo conditions.

To further analyse the interaction and complex behavior in solution, we used AUC, which is an effective tool to understand protein-protein interactions in free solution [24]. Both proteins and complexes in a 1:1 ratio were subjected to ultra-centrifugation and the plotted data suggest the differential mobility in apo and holo conditions (Fig. 1I). In the apo condition, NCS-1 (black) and insulin (gray) showed distinct peaks but the complex displayed a broadly distributed peak against sedimentation coefficient (Fig. 1I). In the presence of 2 mM Ca^2+^, both NCS-1 and insulin showed higher sedimentation coefficient it is shown that Ca^2+^ is part of insulin hexameric structure and stabilises the structure [25]. In the presence of metal ions, the complex spectra line shows multiple peaks that almost coincide with apo insulin, holo insulin, and NCS-1 with another peak of higher sedimentation coefficient. These results suggest that the NCS-1 and insulin forms complexes in apo and holo conditions, which can be observed by AUC.

### NCS-1 binds to insulin

The Trp fluorescence and CD results suggest that insulin interaction induces structural changes in NCS-1 and AUC data indicates the complexes in solution. To further understand the nature of binding we performed titration experiments with ITC. The insulin binding was examined in different conditions like apo, 2 mM Mg^2+,^ and Ca^2+^ conditions. ITC experiments indicate that NCS-1 binds to insulin and the reaction was observed to be exothermic in all experimental conditions [Fig. 2A]. The left image is a representative thermogram for all three experiments done in different conditions and the right panel represents the subtracted and fitted data from different conditions. The NCS-1 binding to insulin is a Ca^2+^ independent event, apo (8.62 µM), in the presence of 2 mM Mg^2+^ (10.5 µM) and Ca^2+^ increases the affinity (1.3 µM) [Fig. 2A and Table 1]. The data was fitted as one set of sites and in all conditions, the N-value was close to 0.5, which suggests one NCS-1 molecule binds to two insulin peptides (NCS-1 was taken in a syringe) (Table 1). The micro-molar affinity of NCS-1 and insulin binding appears to be low considering the extracellular insulin concentration, however, NCS proteins are intracellular and the insulin concentration ranges to moles inside insulin secretory granules, where NCS-1 was shown to localise [1, 6, 26].

**Fig. 2.**
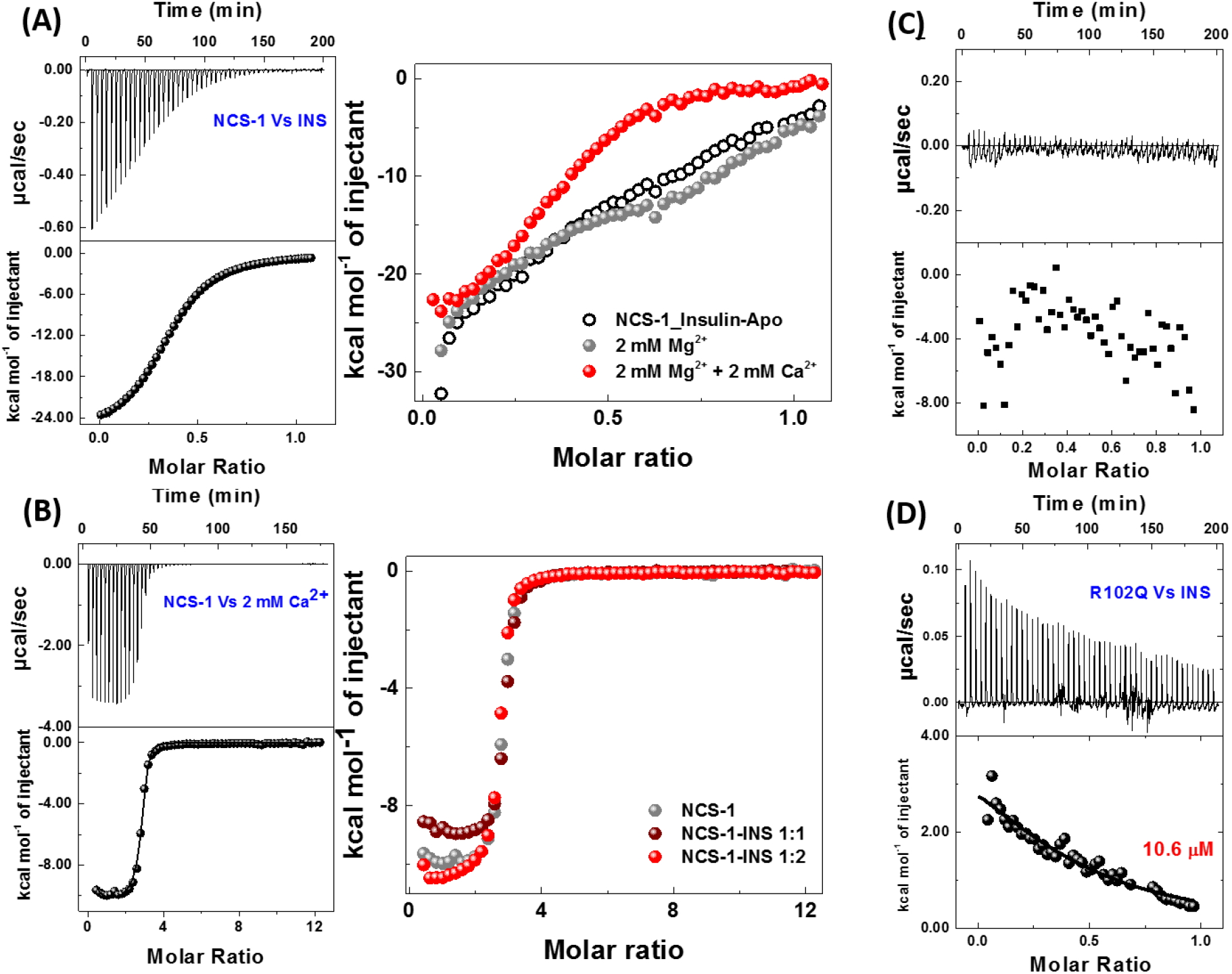
NCS-1 and insulin binding thermodynamics. (A) NCS-1 and insulin interaction by ITC in different conditions apo (black), 2 mM Mg^2+^ (Gray), and holo (Red). Left panel: Representative thermogram and Right panel: Overlap of fitted data from different conditions presented. (B) NCS-1 titration with 2 mM Ca^2+^ in the presence and absence of insulin. (c) ITC representing NCS-1 containing functional 1^st^ EF-hand titration with insulin. (D) NCS-1-R102Q titration with insulin.

**Table 1.**
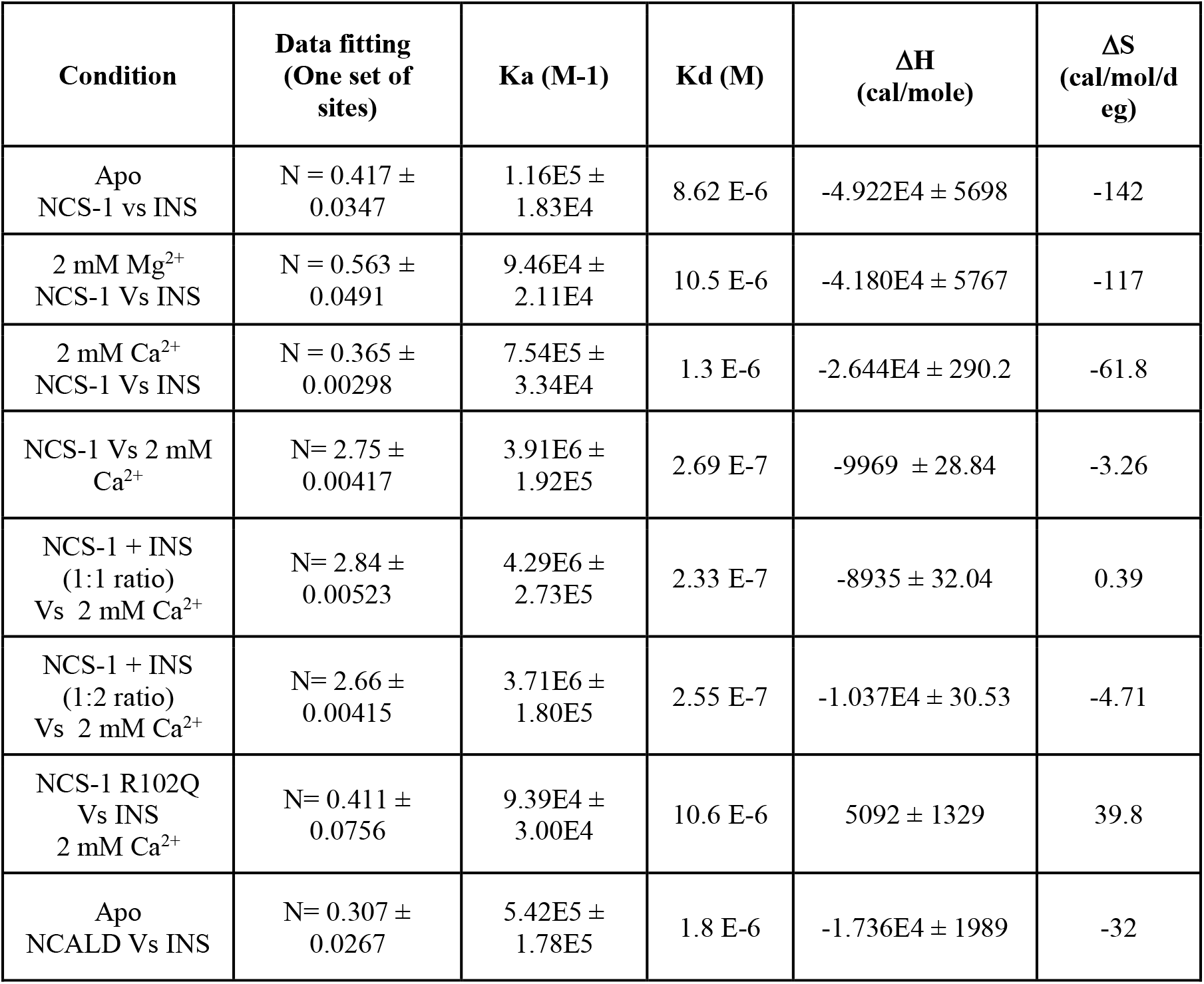
The table represents the thermodynamic parameters obtained from different ITC experiments.

### NCS-1 Ca^2+^ binding properties were unaltered in the presence of insulin

The interacting proteins influence the Ca^2+^ binding properties of sensor proteins, which generate diversity in Ca^2+^ signalling [27]. In an attempt to check phenomena in the NCS-1 system, we examined the effect of insulin-induced structural changes on NVS-1 Ca^2+^ binding properties. The Ca^2+^ titrations were performed on NCS-1 alone, with 1:1 and 1:2 ratios of insulin in the system. In all given conditions the Ca^2+^ binding affinities were unaltered (Fig 2B and Table 1). The left panel is a representative thermogram for all three titrations and the right panel is fitted data plotted as molar ratios. The data suggest that insulin interaction does not alter the Ca^2+^ binding properties.

### Evolutionarily conserved cryptic EF-hand is essential for insulin binding

The NCS proteins are well conserved across species and conserved the first EF-hand as cryptic (non-functional) [1]. NCS-1 conserved 60% of its amino acid sequence and cryptic 1^st^EF-hand from yeast to humans (Fig.S. 1). The mutated NCS-1 protein, where 1^st^ EF-hand made functional (to bind Ca^2+^), is shown to negatively affect the protein’s overall Ca^2+^ binding affinity, conformational flexibility, and activation of PI4Kβ [28]. Here we employed the use of the same NCS-1 mutant to check the effect on the insulin-binding function of NCS-1. The functional 1^st^ EF-hand in NCS-1 resulted in the loss of insulin binding in all experimental conditions i.e apo, Mg^2+,^ and holo (Fig. 2C). The result indicates the functional significance of cryptic EF-hand in NCS-1.

### Single point mutation of NCS-1 alters the insulin binding mechanism

The NCS-1-R102Q mutation identified in autism patient has previously been shown to affect the NCS-1 translocation to the plasma membrane and reduce structural flexibility [29, 30]. Under identical conditions, compared to wild-type, NCS-1-R102Q exhibits a significant change in the mode of insulin binding (Fig. 2A and D). The binding was endothermic in the mutant (in presence of Ca^2+^) and we noticed a substantial reduction in protein expression also. Notably, the incidence of diabetes is also high in autism patients compared to normal individuals [31].

### NCS-1 changes the kinetics of insulin fibril formation

We explored the effect of NCS-1 on insulin fibril formation as insulin can form toxic fibrils and amyloids [20]. The effect of NCS-1 on insulin fibril formation was monitored under apo and holo conditions. NCS-1 affected the rate of fibril formation/nucleation of fibrils compared to insulin alone (Fig. 3A and B). similar lines, the ability of NCS-1 to dissolve preformed fibrils was checked under apo and holo conditions at 60 °C and 25 °C (Fig 3C and D). The results demonstrate that NCS-1 can’t dissolve preformed fibrils.

**Fig. 3.**
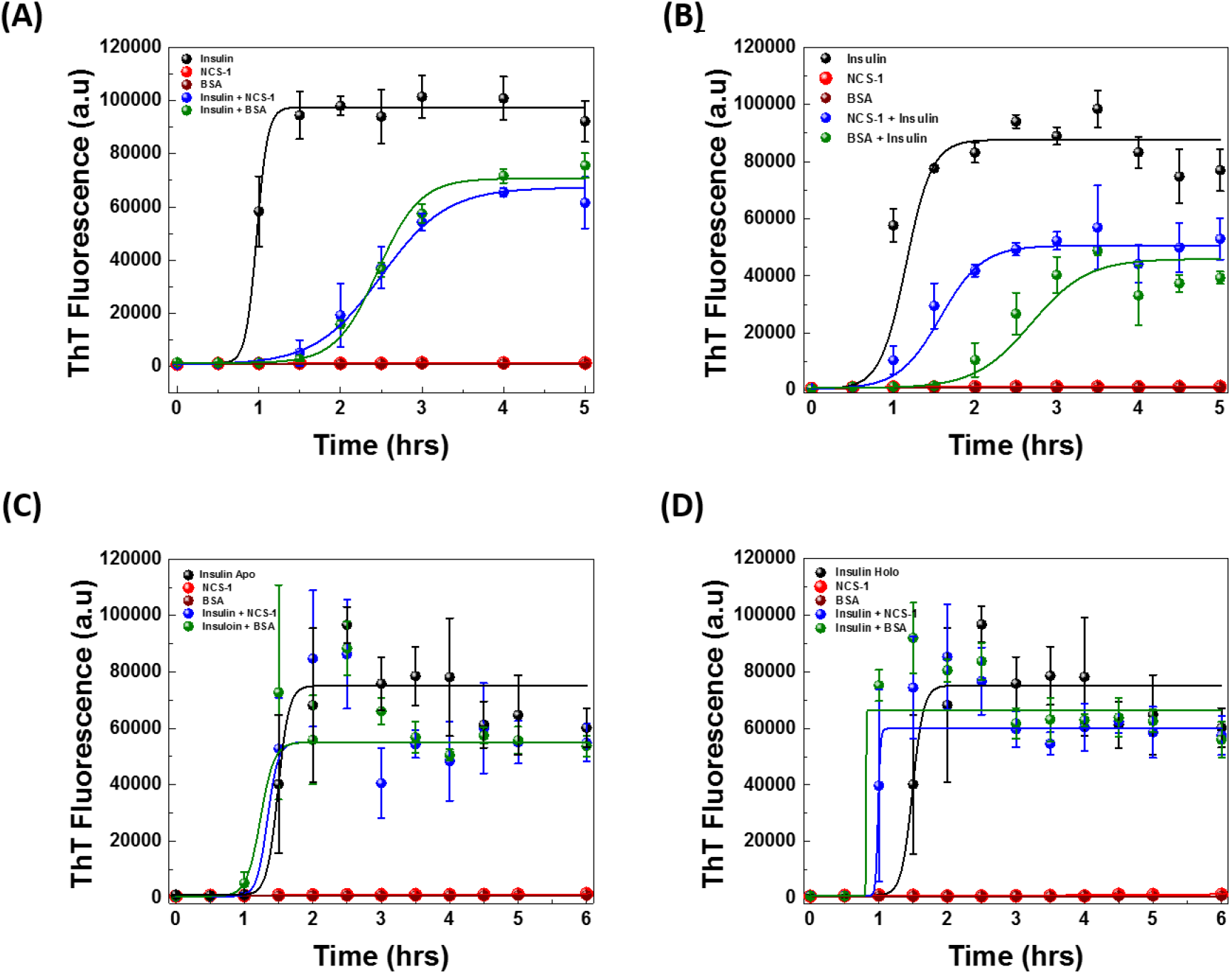
Insulin fibrillation affected by NCS-1. (A) Apo, (B) Holo, the fibrillation kinetics of insulin (Black), NCS-1 (Red), BSA (wine), insulin & NCS-1 (Blue), and insulin + BSA (Green). (C) Apo (D) Holo, reverse fibrillation, presented with an identical colour code.

### Differential regulation of Ca^2+^ on insulin binding of NCS proteins

NCALD is a NCS protein with high similarity to NCS-1 and with limited functional information, expresses in the pancreas [1, 3]. We checked the NCALD interaction with insulin and insulin-induced structural changes by CD and ITC. Results indicate NCALD binds to insulin with an affinity comparable to NCS-1 in a metal ion-independent manner (Fig. 4A, B, and C). Interestingly, Ca^2+^ abolishes the binding despite secondary structural changes shown by CD, which is contrary to NCS-1 (Fig. 4D and F). The insulin-induced changes observed by Trp fluorescence and ANS fluorescence suggest the absence of major structural alterations (Fig.S. 2). The results suggest the differential regulation of Ca^2+^ on the function of NCS proteins. Further, the VILIP was examined for the insulin binding ability in the holo condition only, and the result indicates no detectable binding (Fig. 4F)

**Fig. 4.**
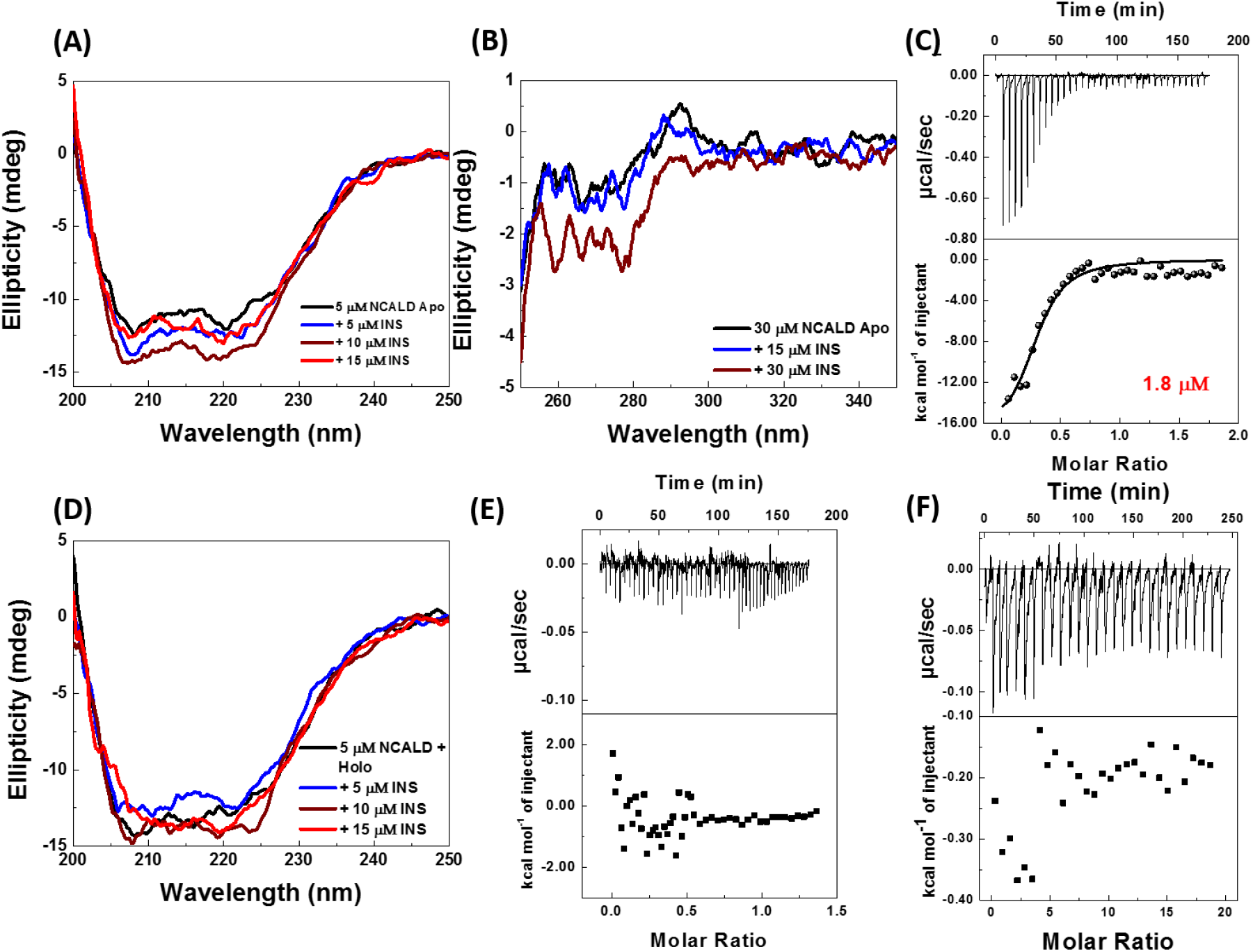
Insulin interacts with NCS proteins. (A) and (B) represents far and near-UV CD spectra of insulin titration with NCALD in apo condition. (C) ITC thermogram of insulin and NCALD titration. (D) Far-UV spectra of insulin titration with NCALD in holo condition. (E) and (F) represents thermograms from insulin titration with NCALD and VILIP respectively in holo condition.

In conclusion, the present work demonstrates that the NCS proteins interact with insulin and Ca^2+^ regulates the binding differently (Fig. 2A and 4E). The expression and significance of different NCS proteins in pancreatic-β-cells were explored previously in the course of exocytosis as they are mainly involved in neuronal secretions (1, 9, 10). In this work, we examined the direct interaction between NCS proteins and insulin through different biophysical applications. The specificity of the insulin interaction with NCS-1 was demonstrated by the significance of conserved cryptic EF-hand and R102Q mutant (Fig. 2C and D). A fibrillation study suggests the possible role of NCS-1 in the stability of insulin in β-cells as NCS-1 is localised to the cytosol, plasma membrane, Golgi, ER, and secretory granules [9]. Insulin and NCS-1 are crucial molecules from embryonic development stages for proper growth and development, hence their interactions at multiple stages must be critical. In addition, the prospective of NCS-1 evolution with insulin physiology can’t be overlooked as insulin-like peptides were identified from *E*.*Coli K*_*12*,_ and glucose tolerance factor was recognized from yeast extract [32, 33]. NCS-1 first appeared in yeast and maintained the cryptic EF-hand throughout, which was shown to essential condition for insulin binding. Another NCS protein NCALD shown to be an insulin-interacting protein and differential Ca^2+^ induced response highlights the new function of NCS proteins to bind insulin. The only shortcoming of this study is we could perform interaction studies of R102Q and VILIP with insulin only in the presence of Ca^2+^. However, insulin synthesis from neuronal progenitors can’t be overlocked, where NCS proteins may have a significant role and these progenitors rescued the diabetic animal from hyperglycemia condition [34]. Further scrutiny of NCS proteins and insulin interaction would help in understanding biological implications and new molecular targets for diabetes treatment.

## Acknowledgment

The authors thank Anand Kumar Sharma and Radhika Khandelwal for providing NCS-1, VILIP, and NCALD clones. We thank Syed Sayeed Abdul for technical help. Thanks to Dr. Santosh Kumar Kuncha for the scientific discussions. J. C Bose National Fellowship (SERB), CSIR, and DST are acknowledged for grants to Y.S. UK recipient of UGC research fellowship.

## Author Contributions

Inception: UK., YS.; Experimental design: UK., Y.S.; Data acquisition: UK., VS; Data analysis: UK., Y.S.; Resources: Y.S.; Writing original draft: UK., Y.S.; Review and editing: UK., Y.S.; Supervision, project administration and funding acquisition: Y.S.

## References

1. Burgoyne, Robert D. “Neuronal calcium sensor proteins: generating diversity in neuronal Ca^2+^ signalling.” Nature Reviews Neuroscience 8, no. 3 (2007): 182–193.

2. Gierke, Paul, Congjian Zhao, Marian Brackmann, Bettina Linke, Uwe Heinemann, and Karl-Heinz Braunewell. “Expression analysis of members of the neuronal calcium sensor protein family: combining bioinformatics and Western blot analysis.” Biochemical and biophysical research communications 323, no. 1 (2004): 38–43.

3. Uhlén, Mathias, Linn Fagerberg Björn, M. Hallström, Cecilia Lindskog, Per Oksvold, Adil Mardinoglu, Åsa Sivertsson et al. “Proteomics. Tissue-based map of the human proteome.” Science (New York, NY) 347, no. 6220 (2015): 1260419–1260419.

4. Boeckel Göran R., and Barbara E. Ehrlich. “NCS-1 is a regulator of calcium signaling in health and disease.” Biochimica et Biophysica Acta (BBA)-Molecular Cell Research 1865, no. 11 (2018): 1660–1667.

5. Grodsky, Gerold M., and Leslie L. Bennett. “Cation requirements for insulin secretion in the isolated perfused pancreas.” Diabetes (1966): 15(12):910.

6. Idevall-Hagren, Olof, and Anders Tengholm. “Metabolic regulation of calcium signaling in beta cells.” In Seminars in Cell & Developmental Biology, vol. 103, pp. 20–30. Academic Press, 2020.

7. Alam, Muhammad Rizwan, Lukas N. Groschner, Warisara Parichatikanond, Liang Kuo, Alexander I. Bondarenko, Rene Rost, Markus Waldeck-Weiermair, Roland Malli, and Wolfgang F. Graier. “Mitochondrial Ca^2+^ uptake 1 (MICU1) and mitochondrial Ca^2+^ uniporter (MCU) contribute to metabolism-secretion coupling in clonal pancreatic β-cells.” Journal of Biological Chemistry 287, no. 41 (2012): 34445–34454.

8. Angebault, Claire, Jérémy Fauconnier, Simone Patergnani, Jennifer Rieusset, Alberto Danese, Corentin A. Affortit, Jolanta Jagodzinska et al. “ER-mitochondria cross-talk is regulated by the Ca^2+^ sensor NCS1 and is impaired in Wolfram syndrome.” Science signaling 11, no. 553 (2018): eaaq1380.

9. Gromada, Jesper, Christina Bark, Kamille Smidt, Alexander M. Efanov, Juliette Janson, Slavena A. Mandic, Dominic-Luc Webb et al. “Neuronal calcium sensor-1 potentiates glucose-dependent exocytosis in pancreatic β cells through activation of phosphatidylinositol 4-kinase β.” Proceedings of the National Academy of Sciences 102, no. 29 (2005): 10303–10308.

10. Dai, Feihan F., Yi Zhang, Youhou Kang, Qinghua Wang, Herbert Y. Gaisano, Karl-Heinz Braunewell, Catherine B. Chan, and Michael B. Wheeler. “The neuronal Ca2+ sensor protein visinin-like protein-1 is expressed in pancreatic islets and regulates insulin secretion.” Journal of biological chemistry 281, no. 31 (2006): 21942–21953.

11. de Rezende, Vitor Bortolo, Daniela Valadão Rosa, Clarissa Martinelli Comim, Luiz Alexandre Viana Magno, Ana Lucia Severo Rodrigues, Paula Vidigal, Andreas Jeromin, João Quevedo, and Marco Aurélio Romano-Silva. “NCS-1 deficiency causes anxiety and depressive-like behavior with impaired non-aversive memory in mice.” Physiology & behavior 130 (2014): 91–98.

12. Ratai, Olga, Joanna Hermainski, Keerthana Ravichandran, and Olaf Pongs. “NCS-1 deficiency is associated with obesity and diabetes type 2 in mice.” Frontiers in Molecular Neuroscience 12 (2019): 78.

13. Mora, Silvia, Paul L. Durham, Jeffery R. Smith, Andrew F. Russo, Andreas Jeromin, and Jeffrey E. Pessin. “NCS-1 inhibits insulin-stimulated GLUT4 translocation in 3T3L1 adipocytes through a phosphatidylinositol 4-kinase-dependent pathway.” Journal of Biological Chemistry 277, no. 30 (2002): 27494–27500.

14. Rossetti, Luciano, Antine E. Stenbit, Wei Chen, Meizhu Hu, Nir Barzilai, Ellen B. Katz, and Maureen J. Charron. “Peripheral but not hepatic insulin resistance in mice with one disrupted allele of the glucose transporter type 4 (GLUT4) gene.” The Journal of clinical investigation 100, no. 7 (1997): 1831–1839.

15. Stenbit, Antine E., Tsu-Shuen Tsao, Jing Li, Rémy Burcelin, David L. Geenen, Stephen M. Factor, Karen Houseknecht, Ellen B. Katz, and Maureen J. Charron. “GLUT4 heterozygous knockout mice develop muscle insulin resistance and diabetes.” Nature medicine 3, no. 10 (1997): 1096–1101.

16. Daneva, Teodora, Shina Pashova, Radoslava Emilova, Plamen Padeshki, Hristo Gagov, and Volodia Georgiev. “DREAM regulates insulin promoter activity through newly identified DRE element.” Open Life Sciences 8, no. 2 (2013): 97–106.

17. Kamiyama, Masumi, Masaaki Kobayashi, Shin-ichi Araki, Aritoshi Iida, Tatsuhiko Tsunoda, Koichi Kawai, Masahito Imanishi et al. “Polymorphisms in the 3′ UTR in the neurocalcin δ gene affect mRNA stability, and confer susceptibility to diabetic nephropathy.” Human genetics 122, no. 3 (2007): 397–407.

18. Okada, Miki, Daisuke Takezawa, Shuji Tachibanaki, Satoru Kawamura, Hiroshi Tokumitsu, and Ryoji Kobayashi. “Neuronal calcium sensor proteins are direct targets of the insulinotropic agent repaglinide.” Biochemical Journal 375, no. 1 (2003): 87–97.

19. Muralidhar, Dasari, Maroor K. Jobby, Kannan Krishnan, Vallabhaneni Annapurna, Kandala VR Chary, Andreas Jeromin, and Yogendra Sharma. “Equilibrium unfolding of neuronal calcium sensor-1: N-terminal myristoylation influences unfolding and reduces protein stiffening in the presence of calcium.” Journal of Biological Chemistry 280, no. 16 (2005): 15569–15578.

20. Sharma, Anand Kumar, Radhika Khandelwal, M. Jerald Mahesh Kumar, N. Sai Ram, Amrutha H. Chidananda, T. Avinash Raj, and Yogendra Sharma. “Secretagogin regulates insulin signaling by direct insulin binding.” Iscience 21 (2019): 736–753.

21. Vivian, James T., and Patrik R. Callis. “Mechanisms of tryptophan fluorescence shifts in proteins.” Biophysical journal 80, no. 5 (2001): 2093–2109.

22. Aravind, Penmatsa, Kousik Chandra, Pasham Parameshwar Reddy, Andreas Jeromin, K. V. R. Chary, and Yogendra Sharma. “Regulatory and structural EF-hand motifs of neuronal calcium sensor-1: Mg^2+^ modulates Ca^2+^ binding, Ca^2+^-induced conformational changes, and equilibrium unfolding transitions.” Journal of molecular biology 376, no. 4 (2008): 1100–1115.

23. Abdurachim, Kholis, and Holly R. Ellis. “Detection of protein-protein interactions in the alkanesulfonate monooxygenase system from Escherichia coli.” Journal of bacteriology 188, no. 23 (2006): 8153–8159.

24. Harding, Stephen E., and Arthur J. Rowe. “Insight into protein–protein interactions from analytical ultracentrifugation.” Biochemical Society Transactions 38, no. 4 (2010): 901–907.

25. Dunn, Michael F. “Zinc–ligand interactions modulate assembly and stability of the insulin hexamer–a review.” Biometals 18, no. 4 (2005): 295–303.

26. Rorsman, Patrik, and Erik Renström. “Insulin granule dynamics in pancreatic beta cells.” Diabetologia 46, no. 8 (2003): 1029–1045.

27. Zhang, Miao, Cameron Abrams, Liping Wang, Anthony Gizzi, Liping He, Ruihe Lin, Yuan Chen, Patrick J. Loll, John M. Pascal, and Ji-fang Zhang. “Structural basis for calmodulin as a dynamic calcium sensor.” Structure 20, no. 5 (2012): 911–923.

28. Rajanikanth, Vangipurapu, Anand Kumar Sharma, Meduri Rajyalakshmi, Kousik Chandra, Kandala VR Chary, and Yogendra Sharma. “Liaison between myristoylation and cryptic EF-hand motif confers Ca^2+^ sensitivity to Neuronal Calcium Sensor-1.” Biochemistry 54, no. 4 (2015): 1111–1122.

29. Handley, Mark TW, Lu-Yun Lian, Lee P. Haynes, and Robert D. Burgoyne. “Structural and functional deficits in a neuronal calcium sensor-1 mutant identified in a case of autistic spectrum disorder.” PloS one 5, no. 5 (2010): e10534.

30. Handley, Mark TW, Lu-Yun Lian, Lee P. Haynes, and Robert D. Burgoyne. “Structural and functional deficits in a neuronal calcium sensor-1 mutant identified in a case of autistic spectrum disorder.” PloS one 5, no. 5 (2010): e10534.

31. Chen, Mu-Hong, Wen-Hsuan Lan, Ju-Wei Hsu, Kai-Lin Huang, Tung-Ping Su, Cheng-Ta Li, Wei-Chen Lin et al. “Risk of developing type 2 diabetes in adolescents and young adults with autism spectrum disorder: a nationwide longitudinal study.” Diabetes care 39, no. 5 (2016): 788–793.

32. LeRoith, Derek, and Lesniak MA. “Insulin or a closely related molecule is native to Escherichia coli.” (1981).

33. Schwarz, Klaus, and Walter Mertz. “A glucose tolerance factor and its differentiation from factor 3.” Archives of Biochemistry 72 (1957): 515–518.

34. Kuwabara, Tomoko, Mohamedi N. Kagalwala, Yasuko Onuma, Yuzuru Ito, Masaki Warashina, Kazuyuki Terashima, Tsukasa Sanosaka, Kinichi Nakashima, Fred H. Gage, and Makoto Asashima. “Insulin biosynthesis in neuronal progenitors derived from adult hippocampus and the olfactory bulb.” EMBO molecular medicine 3, no. 12 (2011): 742–754.

